# GALLANT: A Scalable Computational Platform for Microbial Consortium Interaction and Population Heterogeneity Analysis

**DOI:** 10.64898/2025.12.10.693394

**Authors:** David San León Granado, Juan Nogales

## Abstract

Metabolic modeling enables the prediction of functional capabilities in organisms and microbial communities from genomic data. However, current workflows for genome-scale metabolic model (GEM) reconstruction and contextualization remain time-consuming and technically demanding, particularly when integrating multi-omics data or deriving community-level models from taxonomic profiles. Although recent advances have improved automation and omics integration, challenges persist in incorporating heterogeneous datasets such as single-cell RNA sequencing and in interpreting microbiomes from 16S rRNA data.

We present a computational tool for rapid, automated GEM generation with integrated support for contextualization using transcriptomic and single-cell omics data. The platform also enables the construction of core consortium metabolic models from 16S rRNA profiles, facilitating systems-level interpretation of both single-organism and community-scale datasets. This streamlined pipeline offers a scalable solution for microbiome research, including population heterogeneity analysis and metabolic engineering. Its utility has been demonstrated by exploring phenotypic heterogeneity within Bacillus subtilis populations and identifying metabolic interactions among members of a cyanobacteria-enriched microbial consortium.

**Summary:** Metabolic modeling has become a cornerstone in systems biology, enabling researchers to simulate and analyze the metabolic capabilities of individual organisms and complex microbial communities. Genome-scale metabolic models (GEMs) have been widely used to explore metabolic phenotypes, guide metabolic engineering, and interpret omics data. Despite the increasing availability of genomic and metagenomic data, the generation and contextualization of metabolic models remains a time-consuming and technically demanding process, often requiring manual curation and domain-specific expertise.

At the community level, modeling microbial consortia is gaining momentum, particularly with the rise of microbiome research. Tools like MICOM (Diener et al., 2020) and CarveMe (Machado et al., 2018) have advanced the ability to create metabolic models for entire communities, yet there remains a significant gap when it comes to systematically deriving community-level models from marker-gene data such as 16S rRNA sequencing. While 16S data provides a cost-effective snapshot of microbial composition, translating this into functional, mechanistic models of community metabolism is still an emerging area, typically involving multiple steps of taxonomic inference, genome mapping, and model aggregation.

Recent advances in automation and algorithmic approaches have facilitated the reconstruction of draft models (Tarzi et al., 2024), yet the field continues to face challenges in the accurate contextualization of these models with experimental data. Multi-omics integration, including transcriptomics, proteomics, metabolomics, and especially single-cell RNA sequencing, has the potential to significantly improve the predictive power of metabolic models by providing condition-specific and cell-type-specific constraints. However, integrating these heterogeneous data sources remains a complex task, often limited by compatibility issues, lack of standardized pipelines, and computational overhead.

---

The increasing complexity and scale of bioinformatics analyses, particularly in genome-scale metabolic model (GEM) reconstruction, demand robust, standardized, reproducible, and scalable computational workflows. Workflow management systems such as Snakemake (Mölder et al., 2021) and Nextflow (Di Tommaso et al., 2017) have emerged as essential tools for multi-step pipelines in computational biology (Larsonneur et al., 2018).

In the context of GEM reconstruction, Snakemake enable the automation of complex pipelines, improve reproducibility, and facilitate the parallel processing of multiple genomes or metagenomic datasets. Their use is increasingly critical as metabolic modeling expands to include multi-omics integration, community-scale modeling, and high-throughput comparative analyses.

Its intuitive rule-based structure allows for clear definition of dependencies and execution logic, while its integration with Conda and container technologies ensures reproducible environments across different computing platforms.

To address some of high-level GEM reconstruction limitations, we present a novel tool designed to accelerate and simplify metabolic modeling across scales. Our platform enables the rapid reconstruction of high-quality metabolic models, with built-in modules for integrating omics data, including single-cell sequencing, to contextualize models dynamically. Importantly, it also allows for the automated construction of consensus metabolic models of microbial consortia based on 16S data, inferring a functional “core metabolism” of the community. This approach bridges the gap between taxonomic profiling and systems-level functional analysis, offering a scalable and efficient solution for researchers working with complex microbial datasets.

By streamlining the modeling pipeline and expanding its functional depth, this tool accelerates and simplify metabolic modelling across scales. Our platform enables the rapid reconstruction of high-quality metabolic models, with built-in modules for integrating omics data to contextualize models dynamically. By streamlining the modelling pipeline and expanding its functionality, this tool represents an important advancement in the field, allowing users to do in depth in microbial ecology or synthetic biology.

## Results

### Bioquemistry database for supracellular-scale metabolic modelling

The default database employed is an extended BiGG database. To enhance the predictive capabilities for microbiomes derived from natural environments, where polysaccharide degradation is a prevalent metabolic function, the BiGG database has been extended with other high-quality models reconstructed previously (https://github.com/SBGlab/SBG_GEMs), to include additional reactions involved in the breakdown of complex carbohydrates. Unfortunately, although databases related with exopolysaccharides (EPS) or released polysaccharides (RPS) biosynthesis or degradation are available, e.g., CAZy (Lombard et al., 2014), the information on enzymes, molecular structures and reactions is not assembled together, which makes automated reconstruction of EPS/RPS biosynthetic pathways currently infeasible (Qiu et al., 2023).

Due to the high diversity of EPS/RPS structures (Yoshida et al., 2006), accurately modeling their biosynthetic pathways is highly complex. To reduce this complexity, the variety of cyanobacterial exopolysaccharides was maximally simplified to polysaccharides composed solely of most abundant monosaccharides in cyanobacteria (Laroche, 2022). Additionally, a generic pseudo-metabolite, termed “eps,” was introduced to represent all types of polysaccharides (Table 1). To complete the dataset of common polysaccharides in ecological samples, ulvan, cellulase and alginate degradation enzymes are as well included. The corresponding biosynthetic reactions were assigned to glucosyltransferase genes identified in literature (Pereira et al., 2015), and ABC transport reactions were included whenever the Wzx/Wzy complex genes (Islam and Lam, 2014) were detected.

**Table 1.** List of the main reactions associated to cyanobacterial EPS/RPS biosynthesis and EPS/RPS and algae polysaccharides bacterial degradation.

|  | Reaction | Proteins | References |
| --- | --- | --- | --- |
| **EPS biosynthesis and transporters** | EPSSYN: $\text{glc\_D\_c} + \text{udpg\_c} \rightleftharpoons \text{h\_c} + \text{h2o\_c} + \text{udp\_c} + \text{eps\_c}$<br>EPSabc: $\text{atp\_c} + \text{h2o\_c} + \text{eps\_c} \rightarrow \text{adp\_c} + \text{h\_c} + \text{pi\_c} + \text{eps\_e}$<br>EX_eps_e: $\text{eps\_e} \rightarrow$ | Synechococcus: (Synpcc7942_1144)<br>Synechocystis (slr1534, sll0737, sll5047, slr0728, Slr0977, sll1581) | (Madsen et al., 2023) |
| **Ulvan lyase** | ULVANLY: $\text{ulvan\_c} + \text{h2o\_c} \rightarrow 0.45 \text{ rham\_c} + 0.225 \text{ glcur\_c} + 0.096 \text{ xyl\_D\_c} + 0.095 \text{ glc\_D\_c} + 0.021 \text{ gal\_c}$ | *Algibacter luteus* (SHI30876)<br>*Alteromonas* (WP_120963275, GEA07929.1, WP_052010178, QFR04505.1),<br>*Nonlabens ulvanivorans* (KEZ94292.1),<br>*Pseudoalteromonas agarivorans* (WP_033186995) | (Costa et al., 2022; Gajanayaka et al., 2024) |
| **Alginate lyase** | ALGL: $4.0 \text{ h2o\_p} + \text{prealginate\_M\_p} \rightarrow 5.0 \text{ manur\_p}$ | *Zobellia galactanivorans* (CAZ96774) | (Costa et al., 2022) |
| **Celullase** |  | *Alteromonas* (MG711315.1),<br>*Bacillus amyloliquefaciens* (GU390463.1) | (Barzkar and Sohail, 2020) |

This expansion enables more accurate modelling of microbial metabolic networks, particularly in ecosystems characterized by the abundance of polysaccharide-rich substrates

### Workflow Modularization and description of the main GALLANT characteristics

To optimize computational efficiency, the workflow was modularized by identifying steps that could be executed in parallel due to their independence (Figure 1A). While all steps can be decoupled per model being constructed, a key opportunity for parallelization was found in the reciprocal homology calculation within the core workflow.

**Figure 1.**
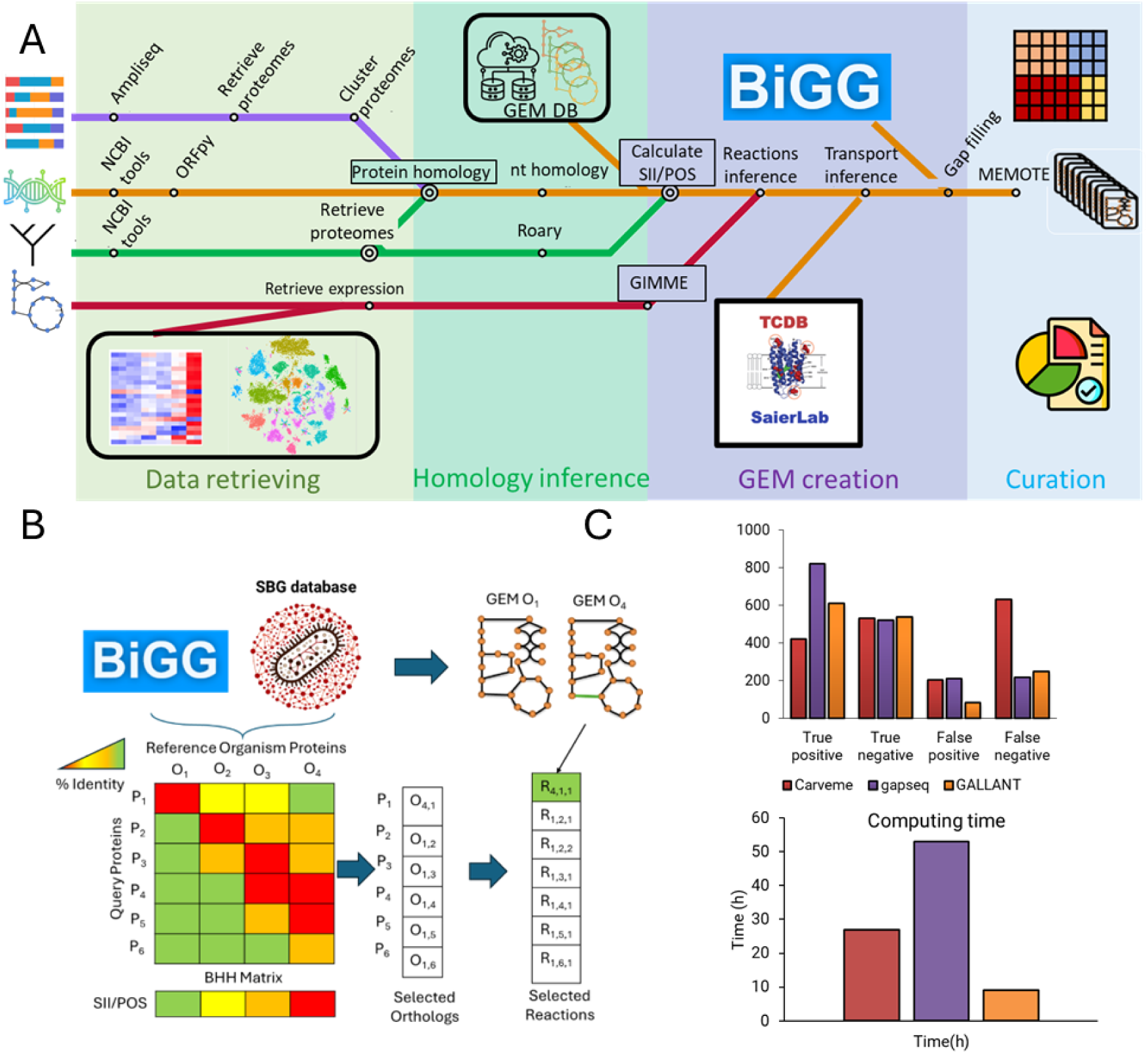
**A**. Schema with the main steps of the extended GALLANT workflow. The orange line is the main pipeline, the purple line represents the sub-workflow to create microbial consortia GEMs workflow to create microbial consortia GEMs from 16S sequencing. The green line represents the sub-workflow to create multi-strain GEMs and the red line represents the sub-workflow to create GEMs from scRNA-seq. **B.** Determination of orthologs and selection of the reactions. The organism proteomes are represented by Oi, i=organism id. The proteins by O i,j, j=protein id, and the reactions Ri,j,k k=reaction id. **C.** Comparison of the results of gapseq, CarveMe and GALLANT. Top) Distribution of TP, TN, FP and FN. Bottom) Computing time of the reconstruction of total ProTraits species database.

Given the distinct characteristics of each step, such as sequence downloading, homology searches, homology searches against the transport sequences, and gap-filling, specific execution rules were defined for each module. This allowed for more effective resource allocation tailored to the computational demands of each task.

Furthermore, each module was encapsulated within a separate Conda environment to prevent conflicts arising from the differing dependencies of the tools employed in each step.

The workflow’s modular design enabled the parallel execution of independent components, significantly improving computational efficiency and scalability. The data retrieval, the sequence annotation if the gene annotation is not provided, the alternative ORF detection and the functional identification are the main steps (Figure 1A).

### A Conservative Strategy for the Functional Identification of Metabolic Orthologs

GALLANT employs a homology-based strategy to identify optimal reaction candidates for metabolic model reconstruction. To mitigate the risk of false positives, the integration of reactions is guided by a global homology proximity criterion: reactions from organisms that are closer to the target species based on total genome homologies are prioritized over those from more distant. Once reactions associated with homologous genes from a given organism are incorporated, the framework restricts the inclusion of additional reactions involving the same gene from other species. This constraint significantly reduces the amount of erroneous functional assignments.

To extract metabolic functional information from a reference database, a strategy inspired by the approach used by OrthoFinder (Emms and Kelly, 2019) was employed (Figure 1A). First, bidirectional best hits (BBH), were identified as the initial criterion for establishing orthologous relationships between genes (Hernández-Salmerón and Moreno-Hagelsieb, 2020).

Next, two complementary metrics were calculated: Average Amino-acid Identity (AAI) (Gerhardt et al., 2025) and Percentage of Orthologs Shared (POS). These metrics allow for the estimation of evolutionary proximity between organisms, which is essential for prioritizing the most functionally relevant orthologous relationships. In particular, the closest organisms were selected, based on the assumption that orthologous genes, especially those involved in central metabolism, tend to have similar functions among closely related species (Altenhoff et al., 2012; Andam and Gogarten, 2011; Bolotin and Hershberg, 2017; Escorcia-Rodríguez et al., 2022). This approach aims to minimize errors in functional assignment.

Protein homology was inferred using the DIAMOND algorithm (Hernández-Salmerón and Moreno-Hagelsieb, 2020), selected for its high speed and accuracy in pairwise comparisons. BLASTp was also available as an alternative. The homology search was divided into three steps to allow further parallelization:

- *Forward mapping*: query sequences were aligned against the reference database.
- *Reverse mapping*: the reference database was aligned against the query sequences.
- *Integration*: results from both mappings were combined to identify bidirectional best hits (BBH).

In cases where a query protein aligned with multiple proteins from another organism, only the best match was retained to ensure specificity.

#### Biomass Reaction Selection Strategy in Genome-Scale Metabolic Model Reconstruction

In the reconstruction of GEMs, GALLANT implements a systematic strategy for biomass reaction selection. Specifically, when no predefined biomass reaction is available for the query organism, GALLANT selects as a template the biomass reaction from the reference organism exhibiting the highest proximity calculated. This choice ensures that the metabolic context is as close as possible to the query organism. To further tailor the biomass reaction, GALLANT automatically adjusts the stoichiometric coefficients of amino acids based on the specific genomic and proteomic composition of the query organism.

Alternatively, if a reference GEM is explicitly provided, GALLANT directly incorporates its biomass reaction, assuming it has been curated and validated for functional relevance.

#### Reaction Transfer

Following the ortholog identification, reactions associated with the orthologous genes were transferred to the target metabolic model. Transport reactions without gene associations were also included. When a query protein mapped to multiple organisms, the selection was based on the organism with the highest proximity, calculated automatically from pairwise comparisons.

#### Transport Reaction Prediction

Transport reactions were predicted based on homology with the curated transporter database (TCDB) (Saier et al., 2021) and the availability of corresponding metabolites in the draft model. These reactions were subsequently integrated into the final metabolic reconstruction.

#### Gap filling

By default, the gap filling is not activated, but if it is activated, the module of gap filling from cobrapy is executed to add the necessary reactions to the model to be able to grow. A special gap filling was developed in the case of very close organism to a specie in the database, for example a different strain. In collaboration with the author of a new gapfilling method (López and Manuel, 2021), a work that focused on improving the automatic generation of genome-scale metabolic models (GEMs) for organisms of relevance in the food industry. The work describes a gap-filling module (GGF) for the GALLANT workflow, for multi-strain GEM reconstruction tool (see Materials and Methods) and shows a clear advantage against classical gap gilling methods. The main objective is to address the limitations of incomplete models resulting from genomes annotated with gaps.

#### Annotation Integration

Annotation of the reconstructed metabolic models is performed at multiple levels to enhance biological interpretability and facilitate downstream analyses.

##### 1. Reaction Annotation

Reactions and metabolites are annotated using information from the BiGG universal model database. If a reaction is present in the BiGG universe, its associated metadata, including reaction identifiers, metabolite names, and stoichiometry, is automatically incorporated into the final model.

##### 2. Gene Annotation

Gene-level annotations can be integrated automatically if the input includes a tabular file containing the following columns: Gene ID, Database, and Database ID. Supported databases include KEGG, NCBI, BRENDA and UniProt. This enables the model to retain links to external biological databases, facilitating cross-referencing and functional analysis.

##### 3. Gene Ontology (GO) Annotation

If a reaction is matched to an entry in the BiGG database, associated Gene Ontology (GO) terms are automatically retrieved and added. This provides functional context to the reactions, supporting enrichment analyses and pathway-level interpretations.

#### Quality Assessment of the Reconstructed Model

The final phase of the workflow is dedicated to assessing the quality, consistency, and completeness of the reconstructed metabolic model. This step ensures that the model meets community standards and is suitable for simulation, analysis, and publication.

As part of this module, the MEMOTE (Metabolic Model Testing) tool is executed. MEMOTE performs multiple tests that evaluate the model across multiple dimensions, including:

- Annotation coverage (genes, reactions, metabolites)
- Stoichiometric consistency
- Mass and charge balance
- Connectivity and network topology
- Presence of blocked reactions
- Compliance with SBML standards

By integrating MEMOTE into the final step of the pipeline, the workflow ensures that models are not only biologically meaningful but also technically robust and reproducible.

### Output Compatibility with External Software

Manual curation of draft metabolic models is one of the most time-consuming yet essential steps in refining model predictions to better reflect biological reality. To support this process and maximize interoperability with a wide range of tools, GALLANT provides model outputs in multiple formats: SBML, MATLAB, JSON, and Excel.

While SBML and MATLAB formats are widely supported by metabolic modeling platforms, the inclusion of an Excel format is particularly remarkable. Given that no existing Python-based tool currently supports full reading and writing of metabolic models in Excel like the version of COBRA in MATLAB, a custom solution was developed specifically for GALLANT, Excel to Cobrapy, a tool that can be accessed in https://github.com/dsanleo/excel_to_cobrapy. This format remains highly valuable, especially for researchers with limited programming experience, as it allows for intuitive inspection, editing, and sharing of models using a familiar spreadsheet interface.

Although the use of Excel in computational biology has deprecated in favor of more standardized formats like SBML, it continues to serve as an accessible and practical option for collaborative model development and dissemination within the scientific community.

#### Carbon source predictions and comparison with other software

One of the potential applications of the generated genome-scale metabolic models is the prediction of carbon source utilization for each organism (Zimmermann et al., 2021). This application was used to evaluate the performance against to other existing tools. To evaluate this capability, we retrieved the ProTraits database, from which we extracted phenotypic data related to growth on various carbon sources. Following the protocol showed by the authors (Zimmermann et al., 2021) the predictions of 1795 carbon sources, 48 carbon sources and 526 organisms, were evaluated with data from the ProTraits database. This dataset was used to assess the predictive accuracy of the models reconstructed for each species present in the database.

We compared the performance of our models with those generated by both gapseq and CarveMe, as these are among the few tools that can be executed in batch mode and do not require commercial licenses or proprietary third-party software, making them suitable for large-scale comparative analyses.

The analysis (Figure 1C) revealed that, although our method achieved lower sensitivity, reflected in a reduced number of true positive predictions relative to gapseq, it demonstrated higher precision, showing a notable reduction in false positive predictions. This suggests a more conservative but potentially more reliable identification of carbon source utilization capabilities.

Regarding execution time of the total model reconstructions (Figure 1C), GALLANT demonstrated a significant performance advantage over both CarveMe and gapseq. The underlying architecture of GALLANT enables faster model reconstruction and batch processing, making it particularly well-suited for large-scale comparative analyses where computational efficiency is critical.

### GALLANT extension to analyze microbial communities and heterogeneity

To enable the modeling of more complex bacterial systems, the core workflow was extended to support a broader range of input data (Figure 1A), thereby expanding its potential applications. Recognizing the diversity of modeling paradigms, top-down, bottom-up, and the hybrid middle-out approach, the workflow was adapted to integrate both and metabolic data from sequenced strains for bottom-up strategies (e.g. multistrain modeling) and metagenomic data (e.g., 16S rRNA sequencing) for top-down designs.

Importantly, it also allows for the automated construction of consensus metabolic models of microbial consortia based on 16S data, inferring a functional “core metabolism” of the community. This approach bridges the gap between taxonomic profiling and systems-level functional analysis, offering a scalable and efficient solution for researchers working with complex microbial datasets.

Nowadays, there is no other tool able to model those supra cellular scenarios (Table 2) and this GALLANT extension fill this lack of functionality.

**Table 2.** Comparison of automatic GEMs reconstruction tools against GALLANT features.

|  | Gap-filling | Default Metabolic database | Strain level reconstruction | Genus-level Metagenomic | Single-cell contextualization |
| --- | --- | --- | --- | --- | --- |
| **Pathway Tools** | Yes | Metacyc | No | No | No |
| **Carveme** | Yes | BiGG | No | No | No |
| **Merlin** | No | KEGG | No | No | No |
| **Raven** | Yes | KEGG | No | No | No |
| **Reconstructor** | Yes | ModelSEED | No | No | No |
| **gapseq** | Yes | ModelSEED | Yes | No | No |
| **GALLANT** | Yes | BiGG | Yes | Yes | Yes |

### Description of workflow

#### Sub-workflow to reconstruct microbial consortia GEMs from 16S sequencing

A specialized sub-workflow was created to reconstruct metabolic models at taxonomic levels beyond individual strains, such as species, genus, or family, based on 16S sequencing data or curated taxonomic lists. This process includes:

- Ampliseq - Initial Processing: 16S RNA amplicon sequencing data (FASTQ format) is preprocessed using the AmpliSeq tool (Straub et al., 2020), which performs quality filtering, taxonomic identification, and abundance estimation. By default, SILVA version 138.1 (Quast et al., 2013) is used as the reference database due to its high-quality curation. This step requires a metadata file following the AmpliSeq input format, and the output will be a phyloseq object (McMurdie and Holmes, 2013).
- Core Microbiome Identification: Assuming that the most abundant taxa contribute most significantly to microbiome functionality, a filtering protocol was implemented using R packages like Microbiome (https://microbiome.github.io/) and Microeco (Liu et al., 2021). Taxa are selected if they represent at least 70% of the total abundance and individually exceed 5% abundance. This approach is particularly suited for engineered consortia in biotechnology, where dominant taxa are expected to drive system behavior. However, it may be less applicable to natural consortia, where taxonomic diversity is high and individual abundances are typically below 1%.
- Retrieve proteomes: For each selected genus, associated proteomes are downloaded with the tool EsMeCaTa (Belcour et al., 2025). EsMeCaTa is a computational method designed to estimate consensus proteomes and metabolic capabilities based on taxonomic affiliations, such as those retrieved from the previous step. It leverages multiple bioinformatics resources, including the UniProt Proteomes database, NCBI Taxonomy or MMseqs2, to generate functional predictions. In this step, EsMeCaTa proteome tool is used to download the proteomes associated to the Core Taxonomy previously generated. If no proteome is available for a given genus, the proteomes of its corresponding family are retrieved as a functional approximation.
- Cluster proteomes: To reduce redundancy and generate a representative reference metaproteome, the retrieved sequences are clustered using EsMeCaTa clustering with default parameters. This step ensures computational efficiency and improves the accuracy of downstream model reconstruction.

After those new computational steps, the rest of the workflow follows the main pipeline, the reaction transference from the metabolic database.

#### Sub-workflow to reconstruct multistrain GEMs

This sub-workflow takes as input a reference strain of a defined specie and search in NCBI database for other full sequenced genomic strains to download them and generate GEMs with the reference model. This workflow reproduced the process implemented at *Bacillus subtilis* multi-strain published previously (Blázquez et al., 2023). The output will be an orthologous matrix, a reactions matrix and the list of metabolic models, one per downloaded genome.

#### Sub-workflow to reconstruct and contextualize microbial GEMs from scRNA-seq

In response to the growing relevance of single-cell sequencing technologies, a dedicated module was also developed to contextualize single-cell RNA-seq experiments, allowing for the reconstruction of metabolic models that reflect cellular heterogeneity.

Specifically, the GIMME algorithm was applied to adapt the generic metabolic model to each cluster’s expression profile, resulting in a set of context-specific models that reflect the metabolic heterogeneity within the bacterial population.

To support this workflow, a new module was developed within GALLANT that accepts single-cell RNA-seq data as input and outputs a collection of metabolic models representing the phenotypic diversity across the population. The module follows the standard preprocessing pipeline defined by the Seurat package in R (Hao et al., 2021). Raw read counts are filtered to retain only genes present in the reference metabolic model. The resulted gene expression was used to generate the list of GEMs with different phenotypes using the GIMME contextualization algorithm. By default, genes are considered active if their expression exceeds the 15th percentile, a threshold chosen to exclude low-expression genes while retaining biologically relevant activity.

This approach enables the reconstruction of metabolic models that reflect the transcriptional heterogeneity of single-cell populations, offering a scalable and biologically informed method for studying metabolic variability at the single-cell level.

### Use case 1: GEM contextualization with single-cell RNA sequencing and heterogeneity analysis

Focusing on *Bacillus subtilis*, contextualizing metabolic models with single-cell RNA sequencing (scRNA-seq) offers a transformative approach to understanding its cellular heterogeneity and dynamic metabolic states. scRNA-seq enables the identification of rare subpopulations and transient physiological states that bulk RNA-seq cannot resolve, which is crucial in a species like *B. subtilis* known for their population heterogeneity under environmental changes (Kearns and Losick, 2005; Kuchina et al., 2021). By capturing gene expression at the single-cell level, it is possible to refine genome-scale metabolic models to reflect real-time cellular behavior, improving predictions of metabolic fluxes and regulatory interactions, enhancing our understanding of how *B. subtilis* adapts to environmental changes. This approach can support applications in synthetic biology or industrial fermentation.

To evaluate the feasibility of integrating single-cell transcriptomic data into GEMs, we utilized a publicly available single-cell RNA sequencing dataset of *Bacillus subtilis* (Kuchina et al., 2021), generated under controlled conditions using LB medium. As a reference, we employed the *i*BB1018 high-quality GEM of *B. subtilis* strain 168 (Blázquez et al., 2023), which served as the reference model for contextualization.

All models were simulated under LB medium conditions to maintain consistency with the experimental setup described by the authors. To assess reaction activity, gene expression data were mapped to their associated reactions, generating what we refer to as reaction expression profiles, which indicate whether a reaction is likely to be active under the given transcriptional state. Expression results were evaluated using a UMAP representation, incorporating OD data mapping (Figure 2A).

**Figure 2.**
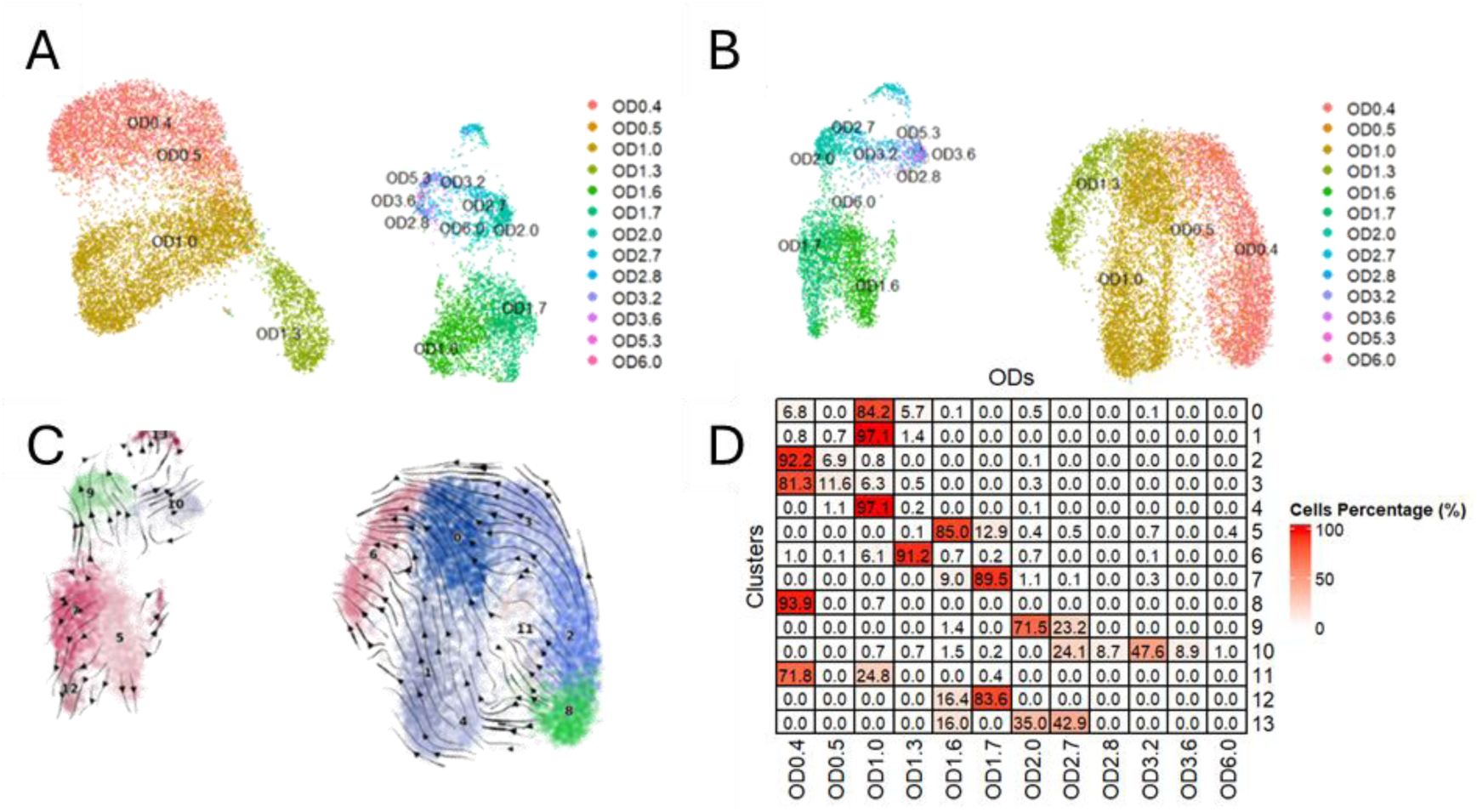
Clustering and UMAP representation of the B. subtilis sc-RNA expression. A. UMAP analysis with the total of gene expression data. B. UMAP analysis with only metabolic genes included into iBB1018 GEM. C. Heatmap with the percentage of shared cells between the calculated clusters (rows) and the ODs (columns). The percentage is calculated against the total of cells per cluster.

To assess the impact of focusing only on metabolic genes with expression data included in our model, we reduced the dataset including only the genes defined in our *i*BB1018 GEM (1018 genes). From them, the total genes with expression in at least one cell was 862 genes (85% of the total genes of *i*BB1018). Later we observed that the overall distribution of cells remained unchanged after dimensionality reduction via principal component analysis (PCA) and UMAP (Figure 2B). This suggests that the metabolic gene subset retains the essential structure of the cellular landscape in this dataset.

To further evaluate whether the data preserved a sense of continuity and directional progression across different optical densities (ODs), we conducted a pseudotime trajectory analysis. The results confirmed a directional flow from early to late ODs, indicating that metabolic changes occur progressively during the development and growth of the microbial community. (Figure 2C)

These findings support the use of GEMs as a robust framework for investigating key metabolic transitions underlying community evolution and adaptation.

After the contextualization of transcriptomic data using the GIMME algorithm, a total of 524 different genome-scale metabolic models (GEMs) were generated, each representing a distinct bacterial phenotype (they are different in at least one active reaction).

Using the GPR integrated into the GEMs, the reaction expression in log scale was calculated. A comparative analysis revealed that several reactions exhibited dynamic expression changes across ODs (Figure 3). For instance, energy-related reactions such as adenylate kinase (ADK1) and deoxyadenylate kinase (DADK) showed a marked decrease in expression during late OD stages, likely reflecting nutrient depletion and reduced energy demand in stationary-phase cells. Furthermore, transport reactions for alternative carbon sources, including cellobiose (CELBpts), arbutin (ARBTpts), and salicin, were upregulated in intermediate ODs, suggesting a metabolic shift toward less-preferred substrates. And in late ODs, upregulation of inositol (inositol (INSTt2). In parallel, reactions associated with glucose utilization (e.g., GLCpts, GLYCt, GLYK, G3PD6) were downregulated, consistent with glucose exhaustion in the medium. Transporters for gluconate (GLYCt) and glutamate (GLUt2r) also showed differential expression, indicating potential shifts in nitrogen and carbon assimilation strategies.

**Figure 3.**
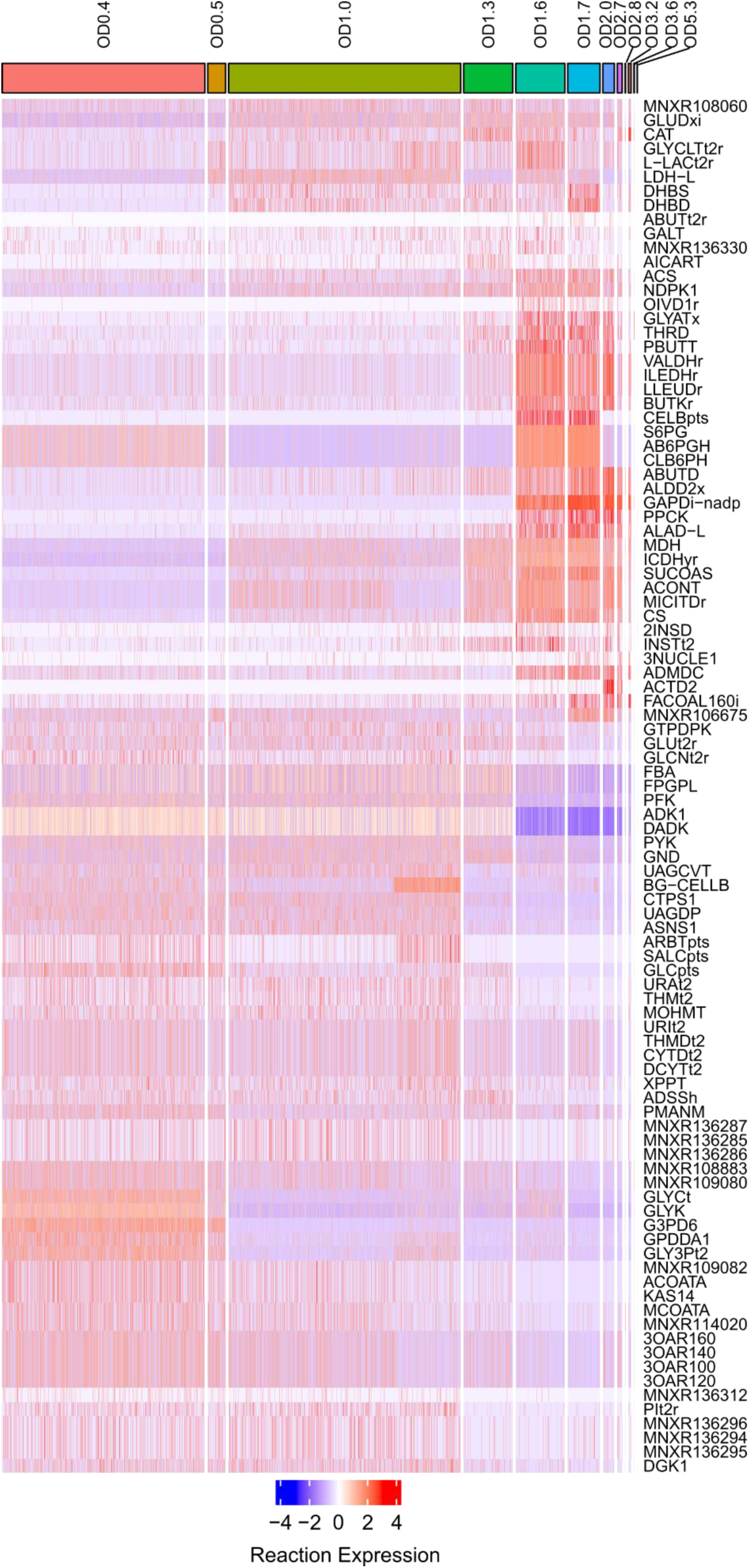
Heatmap wih the log transformed reaction expression of the most differential reactions. Only reactions with GPR are included. The data is row scaled for comparison between ODs.

To minimize errors, we decided to increase the level of expression aggregation until the expression data covered at least the essential genes predicted for *B. subtilis* in LB medium across all clusters.

This approach aimed to reduce potential noise and errors due to sparsity and technical limitations (Nie et al., 2023) from the reduced number of genes with detectable expression per cell in the dataset. By aggregating similar cells, we are able to extract more reliable patterns of metabolic activity and better characterize the phenotypic transitions captured by the GEMs. These approaches enable the detection of biologically subpopulations related with low gene expression coverage. With this approach we identified several phenotypes (clusters) in different ODs. It means that some phenotypes can appear independent of the OD. For example, the cluster 10 represents the phenotype of cells belonging to 9 different ODs.

Following the clustering data processing protocol described by the original authors (Kuchina et al., 2021), and setting the cluster resolution to 2.0, a value, we identified transcriptionally distinct cell clusters characterized by differential expression of metabolic genes (Figure 2D). These clustered data were then used as input for the GALLANT pipeline and 14 GEMs were created.

To analyze the metabolic behavior within distinct transcriptional clusters, median reaction expression profiles were computed across the GEM belonging corresponding clusters, focusing on key metabolic subsystems (Figure 4).

**Figure 4.**
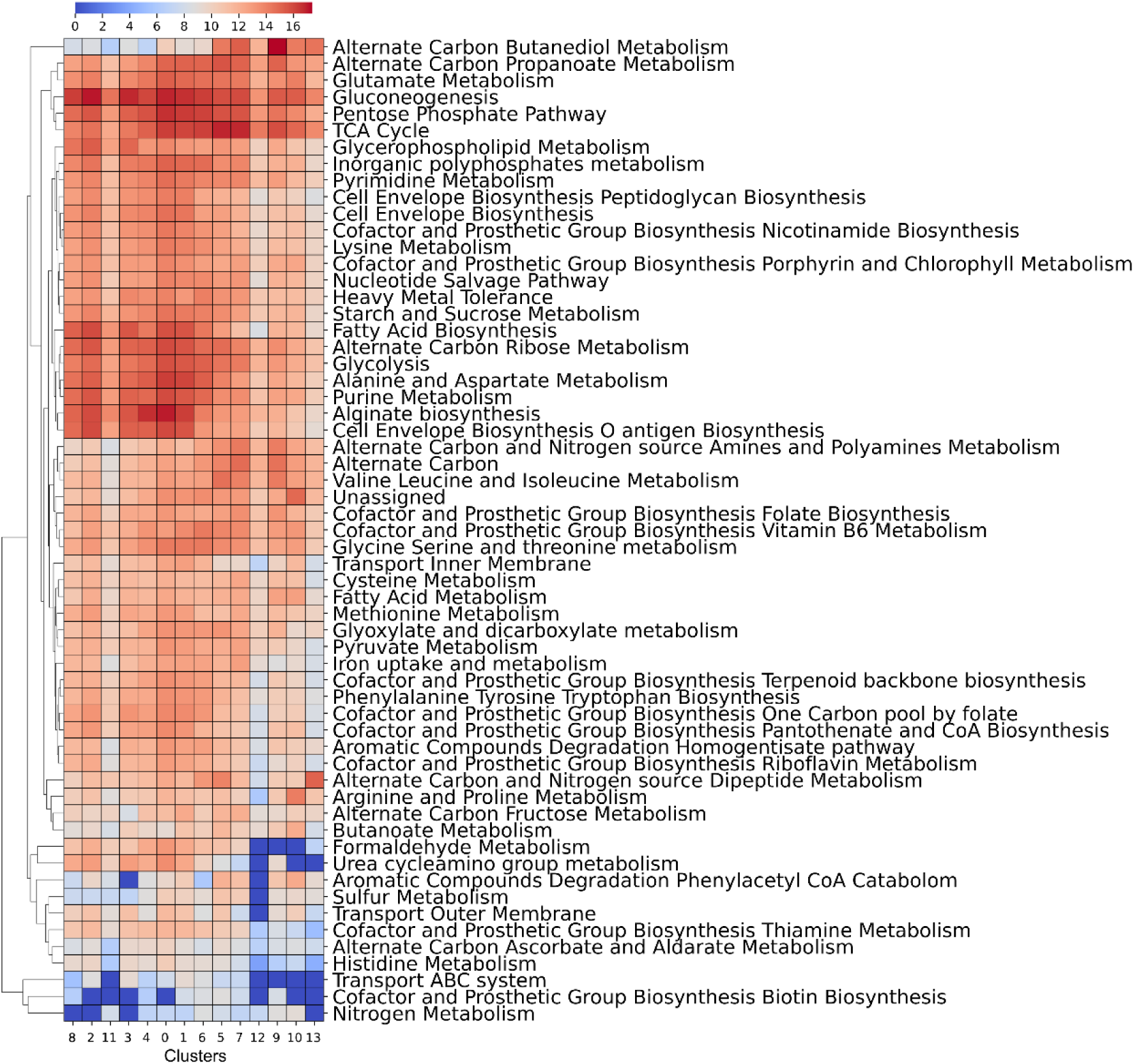
Heatmap with median subsystem expression among the clusters. The order of the clusters is based on the higher percentage of cells for each OD.

Comparative analysis of the resulting models revealed that certain clusters exhibit increased expression through Alternate carbon butanediol metabolism. This suggests a potential metabolic adaptation to localized nutrient depletion experienced by subpopulations of cells. This occur when the flux is redirected from glycolysis to produce 2,3-butanediol and reduce the acetoin or lactate toxic concentration (Yang et al., 2015).

In clusters associated with intermediate ODs (1.0-1.4). The simultaneous upregulation of the tricarboxylic acid (TCA) cycle, pentose phosphate pathway (PPP), gluconeogenesis, glutamate metabolism, and alternate carbon propanoate metabolism reflects a state of high metabolic activity and adaptive flexibility (Blencke et al., 2003; Dergham et al., 2023). This metabolic behavior suggests a shift toward energy-intensive processes, often associated with stress adaptation (Kohlstedt et al., 2014), or nutrient limitation. Gluconeogenesis enables the generation of glucose from non-carbohydrate sources, supporting survival in carbon-limited environments. Upregulated glutamate metabolism contributes to nitrogen assimilation and the synthesis of poly-γ-glutamic acid (γ-PGA), a key polymer in environmental resilience (Sha et al., 2019; Yu et al., 2016). The activation of propanoate metabolism further indicates the utilization of alternative carbon sources, enhancing metabolic versatility (Castillo-Alfonso et al., 2022).

These contextualized models were then subjected to FBA simulations to assess the impact of transcriptional variation on metabolic flux distributions. This approach allowed the identification of subsystems with altered flux capacities, providing a functional link between gene expression heterogeneity and metabolic adaptation (Figure 5)

**Figure 5.**
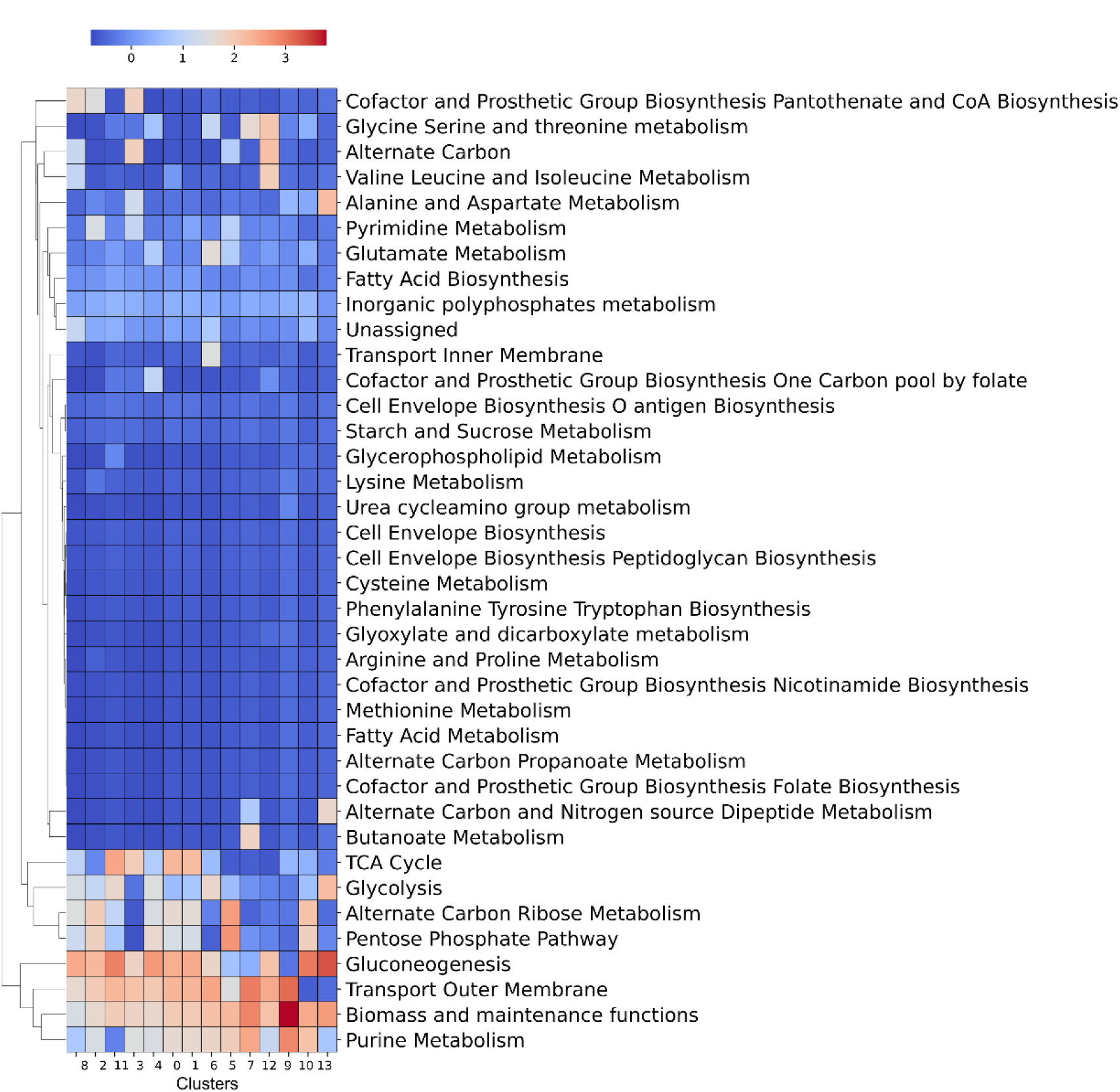
Heatmap with the log absolute median fluxes among the clusters. Only subsystems with flux in at least one cluster are shown.

The flux distributions within the tricarboxylic acid cycle (TCA) varied across clusters (Figure 5), particularly in the medium OD stages of the experiment. These differences may reflect a metabolic shift in response to nutrient exhaustion, supporting the hypothesis that single-cell level heterogeneity can significantly impact metabolic behaviour even within clonal populations.

Other interesting behaviours involve the increase in flux of Alternate Carbon subsystem in different clusters like cluster 10 not associated directly to the OD (Figure 2D). This suggests that individual cells within the same population may adopt distinct metabolic strategies, potentially driven by bet-hedging mechanisms that enhance survival under fluctuating environmental conditions (Grimbergen et al., 2015). It is compatible to the increase of flux of Gluconeogenesis subsystem.

In later growth stages, we observed a significant increase in gluconeogenic flux, anticorrelated with previous expression analysis, which aligns with previous reports linking this shift to the depletion of extracellular carbon sources and the need to maintain anabolic functions (Lim et al., 2024). This metabolic reflects the activation of survival pathways.

A particularly unexpected observation is the upregulation of flux toward alanine and aspartate metabolism in Cluster 13, corresponding to late growth stages. This is counterintuitive, given the central role of these amino acids in core metabolism and their typical downregulation under nutrient stress (Díaz-Pascual et al., 2021).

This behavior could involve cross-feeding mechanisms, where subpopulations secrete metabolites such as alanine that are subsequently utilized by neighboring cells as alternative sources of carbon and nitrogen (Díaz-Pascual et al., 2021). This spatial and functional division of labor has been documented in biofilm communities and may represent a survival strategy in stationary phase. Alternatively, this pattern could be an artifact of modeling, highlighting the need for curation of metabolic models before interpretation.

The analysis of gene expression and metabolic fluxes reveals differences that require further validation. These discrepancies may be due to the absence of key metabolite concentrations in the media at different optical densities (ODs).

Moreover, the assumption that the medium composition remains constant across clusters in LB is a clear limitation for model predictions. It is also possible that higher gene expression does not necessarily correlate with increased metabolic flux.

### Use case 2: Metagenomic modelling- Cyanobacterial Systems for PHB Production

Polyhydroxyalkanoates (PHAs) are a promising alternative for the synthesis of sustainable plastics due to their potential biodegradability, reduced environmental impact and a wide range of applications (Liu et al., 2025) (Jayalath and Alwis, 2025). These biopolymers are naturally produced by bacteria through the fermentation of sugars or lipids (Chien Bong et al., 2021; Kourmentza et al., 2017). However, through metabolic engineering techniques, they can even be synthesized from the degradation of fossil-derived plastics (Tiso et al., 2022) (Liu et al., 2025), offering a potential solution to plastic pollution.

Among the vast diversity of bacteria (Vicente et al., 2023), many studies have focused on PHA production using heterotrophic bacteria (Manoli et al., 2024), (Inoue et al., 2016). An interesting alternative is the use of cyanobacteria, which, being autotrophic, can accumulate PHB using sunlight and CO₂ as primary resources. This opens the possibility of producing polymers within a circular economy framework, removing greenhouse gases from the environment. Cyanobacteria accumulate PHB under nutrient-limited conditions as a strategy for resource storage. However, PHB production in cyanobacteria is generally lower than in heterotrophic bacteria, prompting the development of new strategies to enhance yields (Altamira-Algarra et al., 2025, 2023). Since pure cultures are often inefficient, natural microbial consortia enriched in cyanobacteria have been employed. These consortia offer advantages such as eliminating the need for sterilization and expanding the range of usable carbon sources (Altamira-Algarra et al., 2025).

To test the GALLANT functionalities with top-down engineering approaches and to investigate hypotheses derived from metagenomic studies, we analyzed publicly available metagenomic datasets. In particular, we focused on the dataset published by Altamira-Algarra et al. (Altamira-Algarra et al., 2025), which employed two cyanobacteria-enriched consortia for polyhydroxybutyrate (PHB) production. This study implemented a two-phase cultivation strategy, previously demonstrated to be effective for sustained PHB production in cyanobacteria-dominated microbiomes (Altamira-Algarra et al., 2023). Our objective was to identify key differences in microbial interactions within consortia exhibiting high PHB productivity.

To this goal, we analyzed the metabolic potential of the most abundant genera, aiming to determine whether their functional capabilities, along with the heterotroph-to-autotroph ratio, could explain the observed differences in PHB production. As a starting point, we selected the stabilized microbiomes CW1 and CW2 at day 160 (last Cycle) of the experiment. The core consortium of each microbial community, defined as the set of most abundant taxa (bigger than 2% of relative abundance in at least one sample), was identified using the GALLANT tool. These core taxa are presumed to be the primary contributors to the community’s functional output. There are no differences in the genera detected between the two microbiomes, but differences in relative abundance are evident.

The generated consortium is primarily composed of a cyanobacteria, an alphaproteobacteria, and a gammaproteobacterial, a configuration previously described in cyanobacterial consortia (Zheng et al., 2020). Based on the identification of key genera such as *Synechocystis*, *Pseudoxanthobacter, Methyloversatilis* and *Mesorhizobium* (Table 3), we used GALLANT to retrieve the unified proteomes of organisms with available sequences in UniProt, selecting a single representative per homologous protein group. The abundance of the selected samples was almost the 90% of the total abundance, so it can be a good presentative microbiome with a reduction of complexity. The resulting dataset constitutes a genus-level proteome, which was used to construct genus-level metabolic models. Only 3 models were possible to generate due to the lack of public complete genomes in NCBI/Uniprot databases of *Mesorhizobium* genus, the less abundant genus of the dataset.

**Table 3.** List of the genus included into the selected core-microbiomes with their relative abundance (%). The microbiome with higher PHB production (CW1) and the microbiome with lower PHB production (CW2).

| Class | Genus/Specie | CW1 | CW2 | Number of Genomes |
| --- | --- | --- | --- | --- |
| *Cyanobacteria* | *Synechocystis PCC-6803* | 48.88 | 63.81 | 2 |
| *Alphaproteobacteria* | *Pseudoxanthobacter* | 36.3 | 22.89 | 3 |
| *Gammaproteobacteria* | *Methyloversatilis* | 2.59 | 2.11 | 25 |
| *Alphaproteobacteria* | *Mesorhizobium* | 2.12 | 0.55 | 0 |
|  | **Total relative abundance (%)** | 89.92 | 89.36 |  |

Subsequent analysis of the constructed models revealed that the main autotrophic specie, *Synechocystis*, produce polysaccharides that are subsequently can be utilized by other community members, a well-known behaviour (Pan et al., 2023), which degrade these compounds and use them as carbon sources. Inoculum ratios were constrained based on 16S rRNA relative abundance data and the selected biomass (400 mg/L). The maximum microbiome growth was constrained with the detected by the authors (0.05 d^-1^).

This polysaccharide-mediated interaction plays a crucial role in maintaining the carbon cycle within the consortium. Using dynamic flux balance analysis (dFBA), it was observed that the system was not stable with the initial produced models. With photoautotrophic *Synechocystis* metabolism, *Pseudoxanthobacter* is not able to grow (Figure 6A) something that it does not fit with the experimental results and the genus *Methyloversatilis* is able to grow with the remaining acetate and citrate as carbon source.

**Figure 6.**
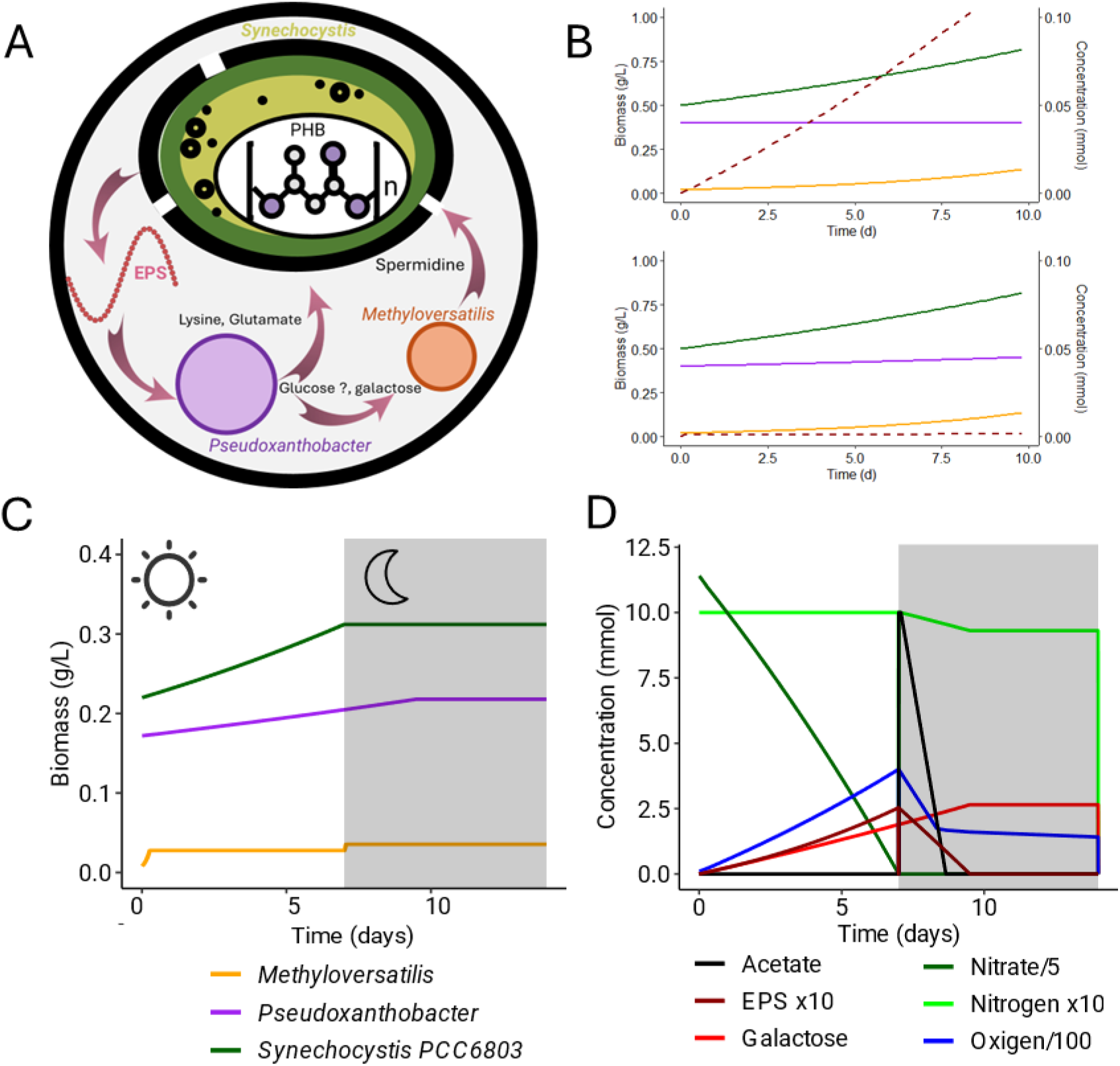
**A**. Schema of the predicted interactions of the microbiomes. **B**. COMETS simulation of Synechocystis (green), Pseudoxanthobacter (purple) and Methyloversatilis (orange) and EPS/RPS accumulation (dotted line) Top) Microbiome without EPS/RPS degradation functionality, Bottom) With EPS/RPS degradation. **C.** dFBA simulation of a total cycle of the microbial consortia. Left) Biomass abundances. Right) Metabolite concentrations. Some of metabolite concentrations have been scaled to make the figure easier to interpret.

Consequently, to test if the reason is a missing polysaccharide degradation functionality, a detailed evaluation of the consensus proteome of *Pseudoxanthobacter* was conducted against the CAZy (Carbohydrate-Active enZYmes) database (Lombard et al., 2014) using the tool dbSCAN3 server (Zheng et al., 2023) to identify potential enzymes involved in the utilization of alternative carbon sources beyond those predicted by the initial metabolic reconstruction. This analysis revealed the presence of several enzymes belonging to glycoside hydrolase families (Table 4), which are known for their role in polysaccharide degradation. A total of 35 Glycoside Hydrolases were detected and seven of these glycoside hydrolases contained predicted signal peptides, suggesting their potential for extracellular secretion. Interestedly, several types of substrates were predicted and some of them related to polysaccharides like starch or cellulose.

**Table 4.** List of glycoside hidrolases detected by scandb against CAZy database.

| GLYCOSIDE HYDROLASES |  | NUMBER OF ENZYMES |
| --- | --- | --- |
| **TOTAL** |  | 35 (7 with signal peptide) |
| **PREDICTED SUSTRATES** | Starch | 5 |
|  | Sucrose | 7 |
|  | Glycogen | 5 |
|  | Chitin, Xylan | 5 |
|  | Cellulose | 4 |
|  | Beta-glucan | 3 |
|  | Unknown | 6 |

**Table 5.** Results of main potential cross-feeding metabolite exchange in growth phase.

| Metabolite | Produced by | Consumed by |
| --- | --- | --- |
| Pyruvate | *Synechocystis* | *Methyloversatilis* |
| 2-Oxoglutarate | *Synechocystis* | *Pseudoxanthobacter* |
| Oxygen (O<sub>2</sub>) | *Synechocystis* | *Methyloversatilis*, *Pseudoxanthobacter* |
| EPS/RPS | *Synechocystis* | *Pseudoxanthobacter* |
| L-Lysine | *Pseudoxanthobacter* | *Synechocystis*, *Methyloversatilis* |
| H<sub>2</sub>S | *Pseudoxanthobacter* | *Methyloversatilis* |
| Indole | *Pseudoxanthobacter* | *Methyloversatilis* |
| Acetate | *Methyloversatilis* | *Synechocystis* |
| Ammonium | *Methyloversatilis* | *Synechocystis*, *Pseudoxanthobacter* |
| CO<sub>2</sub> | *Methyloversatilis*,<br>*Pseudoxanthobacter* | *Synechocystis* |
| L-Glutamate | *Pseudoxanthobacter* | *Synechocystis*, *Methyloversatilis* |
| Spermidine | *Methyloversatilis* | *Synechocystis* |

The results could indicate that this genus requires polysaccharide degradation abilities to use EPS/RPS as carbon source. Unfortunately, the lack of annotated genes for polysaccharide degradation in our metabolic database avoided the prediction.

To further explore system dynamics using dynamic Flux Balance Analysis (dFBA), a pseudo-reaction (Equation 1) was incorporated into the *Pseudoxanthobacter* genus level metabolic model to simulate the degradation of EPS into oligosaccharides. This modification allowed the model to account for EPS/RPS turnover and nutrient recycling, leading to a more stable and realistic simulation of microbial behavior under dynamic conditions, consistent with previous approaches in microbial systems modeling. (Figure 6B)

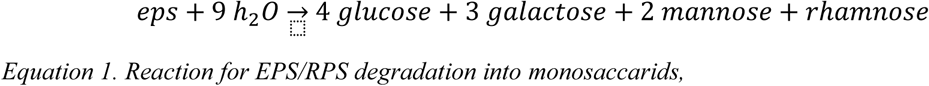

To test potential interactions under not optimal optimization, the genus-level GEMS were aggregated into a community GEM. The individual models were integrated into a separated artificial compartment and a new comparted was created to simulate the medium to use it as inter-specific exchange medium. Additionally, potential metabolic exchange relationships within the microbial consortium were identified using community FVA as an approach to identify possible exchanges in not optimal solution. In the results, it was observed that *Pseudoxanthobacter* can produce some aminoacids like lysine and glutamic acid that can be consumed by *Synechocystis*.

Other interesting metabolite, spermidine is a polyamine known to play a role in stress response, environmental adaptation and cellular signaling in cyanobacteria (Zhu et al., 2015), (Jantaro et al., 2014; Yodsang et al., 2014), the same with the close related polyamine, putrescine (Raksajit et al., 2006). It suggests a potential regulatory mechanism influencing glycogen/PHB biosynthesis due to the stress conditions.

Related to the *Methyloversatilis* and *Pseudoxanthobacter* potential for polyhydroxyalkanoate (PHA) or polyhydroxybutyrate (PHB) accumulation remains uncertain, since no associated biosynthetic genes have been identified in their proteomes and associated GEMs.

To see the behavior of the system in dark conditions, the dFBA simulation was extended in time, the light was removed from the *in silico* medium and acetate was added as carbon source (10 mmol).

With this new situation, *Synechocystis* and *Methyloversatilis* were not able to grow due to the nitrogen limitation, but *Pseudoxanthobacter* is able to grow (Figure 6C & D) while EPS/RPS is available.

The *Pseudoxanthobacter* model has as well the ability to assimilate nitrate and it has the presence of a nitrogenase enzyme, which enables the fixation of atmospheric nitrogen (N₂) into bioavailable ammonium (NH₄), a capability to grow under nitrogen starvation. In general, the nitrogenase activity is inactivated by oxygen presence but in some cases like *Azotobacter vinelandii* (Alleman et al., 2021) this activity is maintained even with oxygen, so this result should be confirmed experimentally.

As well, it was detected that a proportion of sugars like galactose (Figure 6D) are accumulated into the medium due to the *Pseudoxanthobacter* degradation process. The presence of these sugar, together with residual acetate, may enable *Synechocystis* to adopt a photomixotrophic metabolic state, in which light-driven photosynthesis is complemented by the assimilation of organic carbon sources. This metabolic flexibility can enhance glycogen accumulation during the growth phase. As a result, during subsequent nutrient starvation, the increased glycogen reserves may lead to higher polyhydroxybutyrate (PHB) production, improving the overall yield of bioplastic synthesis (Dutt and Srivastava, 2018).

The predicted interactions (Figure 6A**¡Error! No se encuentra el origen de la referencia.**) don’t just point to possible cross-feeding or resource sharing. EPS/RPS are key in microbial biofilms, especially those formed by cyanobacteria. They provide structure and store carbon. If *Pseudoxanthobacter* can degrade these polymers, it could help recycle carbon, which might affect nutrient access and community dynamics. Its potential ability to fix nitrogen adds another layer: it could support the ecosystem when nutrients are limited, complementing cyanobacteria and helping the whole community stay balanced. Plus, the predicted cooperation with *Methyloversatilis* under stress, through spermidine exchange, suggests a stabilizing effect that could protect the consortium from environmental shocks. These ideas come from GEM-based simulations, so the next step is to test them experimentally to see if they hold true. These findings show the importance of integrating genome-scale metabolic modeling with community-level analyses to uncover emergent properties in microbial consortia.

## Materials and Methods

### GALLANT Workflow

The development of GALLANT was implemented using Snakemake (version 9.14.4), a workflow management system designed for reproducible and scalable data analyses. To ensure robust dependency management and avoid conflicts between software packages, each rule and module within the workflow was encapsulated in its own isolated Conda environment (version 25.9.1).

#### Individual modules

##### Data Retrieval

Genomic and proteomic sequences were retrieved using NCBI Datasets CLI Tools v 18.0.5 based on provided NCBI identifiers. In cases where custom sequences were supplied, this step was omitted.

##### Sequence Annotation

When annotations were not available, functional inference was performed using the EggNOG tool (Cantalapiedra et al., 2021), allowing the pipeline to proceed with downstream analyses.

##### Alternative ORF Detection

Alternative open reading frames (ORFs) were identified using the orfipy (Singh and Wurtele, 2021) tool with the following set parameters --min 10 --max 10000, ensuring comprehensive coverage of potential coding regions.

##### Homology inference. Diamond parameters

Default thresholds were set as follows: e-value < 0.05, identity > 30%, and minimum coverage ≥ 40%. These parameters were configurable via the workflow’s settings file.

##### Transport reactions

The TransportDB database (Saier et al., 2021) was used as transport gene database to infer transport reactions for the metabolic reconstruction.

### Supra cellular metabolic model reconstructions and analysis

#### B. subtilis Single-cell model reconstruction and analysis

The single-cell RNA sequencing data from B. subtilis were analyzed using the R package Seurat (version 5). The clustering was done with the function FindClusters with the resolution parameter set to 2.0. Gene expression for the different groups of cells sharing the same phenotype was calculated using the AggregateExpression function. These expression profiles were then used to compute the genome-scale metabolic models (GEMs), which were generated with the GIMME algorithm integrated into GALLANT with a threadshold set to 0.15.

#### Cyanobacterial-riched SMC reconstruction and analysis

The microbial consortia GEM was assembled with the pyCOMO framework [51], with the fixed growth option. Potential metabolic exchange relationships within the microbial consortium were identified using community Flux Variability Analysis (FVA), implemented in the pyCOMO framework [51] with default configuration. The dFBA simulations were performed with COMETS 2 software [50]. The flux sampling analysis was run with optgp method implemented in COBRApy generating 10000 samples by run. The dFBA analysis was driven by COMETS v2.

#### Computational runtime

The runtime was measured on a system running Linux Ubuntu 20.04 with 50 GB RAM and an Intel(R) Xeon(R) Gold 6230R CPU @ 2.10GHz using 48 cores.

## Discussion

GALLANT introduces a modular, scalable workflow for genome-scale metabolic model (GEM) reconstruction, effectively overcoming key limitations of current methodologies. Its flexible architecture enables seamless integration of new modules, fostering continuous enhancements in model accuracy, annotation depth, and functional coverage.

Beyond core metabolic pathways, GALLANT extends its scope by incorporating a curated dataset of reactions and genes involved in polysaccharide synthesis and degradation. This addition improves the representation of ecologically relevant interactions, such as those observed in cyanobacteria–heterotroph consortia and marine microbiomes associated with algal communities.

In contrast to existing tools like CarveMe and gapseq, GALLANT delivers high-throughput GEM generation with consistency and efficiency, while preserving biological relevance and contextual fidelity.

### Contextualization of a single cell GEMs of B. subtilis

A large set of genome-scale metabolic models (GEMs) representing different phenotypes within a monoclonal *Bacillus subtilis* community was generated across various optical densities (ODs) during growth. Although the limited number of genes with expression data, due to technical constraints, prevents accurate flux predictions at the scale of a few cells, the approach proved useful for assessing the expression of genes associated with metabolic reactions. The analysis revealed potential differences in metabolite consumption across growth stages, opening the possibility of predicting metabolite concentrations over time or in previously uncharacterized media.

To address the limitations caused by sparse expression data, clustering resolution was reduced to improve reliability, albeit at the cost of losing some phenotypic variability. Among the identified phenotypes, one exhibited an increase in the gluconeogenesis subsystem, which is consistent with bet-hedging mechanisms that enhance survival under fluctuating environmental conditions (Grimbergen et al., 2015). Additionally, an upregulation of flux toward alanine and aspartate was observed at late ODs, where resources are limited, suggesting potential cross-feeding interactions similar to those reported in other organisms. These findings indicate that single-cell level modeling enables the generated models to capture internal interactions among individual cells, such as division of labor (DoL) strategies.

### Microbial consortia modeling following top-down engineering process

A genome-scale metabolic model (GEM) of a microbial consortium enriched in PHB-accumulating cyanobacteria was constructed. This reconstruction enabled the identification of metabolic gaps, notably the potential degradation of cyanobacterial EPS/RPS by members of the genus *Pseudoxanthobacter*. Such findings underscore the capacity of GEM-based approaches not only to identify known interspecific interactions but also to infer previously undescribed relationships or metabolic functions that remain hidden due to limitations in gene annotation. This predictive capability is particularly relevant for uncovering ecological roles and metabolic complementarities within complex microbial communities and this can be improved in the future following strategies like community gap-filling (Giannari et al., 2021).

Beyond gap identification, the model facilitated the evaluation of potential metabolite exchanges among consortium members. These analyses revealed key interactions that may enhance glycogen accumulation in *Synechocystis* and contribute to its stabilization. For instance, the exchange of glutamate produced by *Pseudoxanthobacter* can allow higher flux to glycogen synthesis by *Synechocystis*. Spermidine produced by *Methyloversatilis* suggests a cooperative metabolic network that supports cyanobacterial energy storage under consortium conditions. Such interactions could represent adaptive strategies that improve resilience and resource allocation within the community.

The analysis employed Flux Variability Analysis (FVA) implemented in PyCOMO (Predl et al., 2024) as an initial approach to approximate cross-feeding interactions, providing a basic analysis for studying potential metabolic exchanges between species. While this method offers a useful starting point, it is relatively limited in its ability to capture the complexity of ecological relationships. More sophisticated tools such as MICOM (Diener et al., 2020) or COMMA (Mirzaei and Tefagh, 2024), which implement multiple community-level modeling strategies, could significantly refine these interaction predictions. These platforms enable the identification of specific ecological dynamics, including competition, mutualism, and parasitism, thereby offering a better understanding of microbial consortia. This information could enhance the COMETS simulations and explore how metabolic exchanges and ecological relationships evolve dynamically in heterogeneous environments.

Nevertheless, it is essential to emphasize that these predictions require experimental validation. The modeling process aggregates metabolic functions across all members of a given genus, which may overestimate or generalize capabilities that are strain-specific. Furthermore, the reliance on curated annotations introduces inherent limitations.

### Future GALLANT extensions

In the current state of the art, very few tools combine GALLANT functionalities into a single platform, although several implement them individually. For example, metaGEM (Zorrilla et al., 2021) generates consortium-level models from metagenomic data, relying on CarveMe for reconstruction. Similarly, panDraft (De Bernardini et al., 2024), part of the gapseq framework (Zimmermann et al., 2021), can reconstruct complete metabolic models at the species level for uncultivable organisms using metagenomic assemblies. This tool addresses incomplete genomes by clustering sequences from the input dataset to create a composite genome for modeling purposes.

GALLANT, in contrast, currently it is restricted to complete genomes to minimize errors and ensure higher accuracy. However, the approach used by panDraft is conceptually compatible with GALLANT’s extension toward genus-level metabolic models, as it involves merging proteomes from both complete and incomplete genomes to form a unified representation. Incorporating such functionality into GALLANT would be relatively easy and could significantly increase its applicability to specie-level contexts.

GALLANT’s architecture enables the possibility of integration of new modules, which not only improves model quality, annotation depth, and functional scope but also lays the foundation for future enhancements. Thanks to this modularity and scalability, incorporating additional capabilities, such as advanced annotation tools, improved gap-filling strategies, or context-specific metabolic constraints, will further reduce current limitations and expand GALLANT’s applicability in diverse research and industrial settings. For example: Reproducibility Reporting: Integration of FROG reports to guarantee numerical reproducibility and streamline model curation, leveraging prior experience with FROG publication (Raman et al., 2024).

GALLANT is still under development and presents certain limitations, primarily due to its dependence on a relatively small and curated metabolic database compared to larger, well-established resources such as KEGG (Ogata et al., 1999) or MetaCyc (Caspi et al., 2020). While it is true that the size of GALLANT’s database can grow as new genome-scale reconstructions become available, this expansion remains constrained by the completeness and accuracy of the underlying data. One potential strategy for overcoming this limitation in the future would be to incorporate these larger databases as secondary sources of metabolic information. However, to preserve one of GALLANT’s key advantages, minimizing false positives, reactions imported from secondary databases should be assigned lower priority relative to those from the primary curated database. This hierarchical approach could help maintain the balance between comprehensiveness and reliability. Another promising avenue is the development of a large-scale metabolic network for gap-filling purposes, or the integration of predicted metabolic networks generated by tools such as AtlasX (MohammadiPeyhani et al., 2022). These enhancements could significantly improve the coverage and accuracy of metabolic reconstructions, particularly for poorly characterized organisms.

These new functionalities open the door to modeling large natural consortia, where calculating the metabolic potential of diverse associated genera can progressively refine our understanding of their ecological interactions. Using GALLANT’s automated capabilities, it would be feasible to use the complete NCBI taxonomy as a starting point to develop a comprehensive database of genome-scale metabolic models (GEMs) at the genus level. Such a resource would enable systematic evaluation of metabolic potentials and functionalities, allowing researchers to cluster organisms and build a portfolio of candidate genus or species for the design of synthetic microbial communities (SMCs) through middle-out engineering strategies.

## Author Contributions

Conceptualization, D.S.L. and J.N.; methodology D.S.L. and J.N.; software, D.S.L.; formal analysis, D.S.L.; data curation, D.S.L. and J.N.; writing—original draft preparation, D.S.L. and J.N.; writing—review and editing, D.S.L., J.N.; supervision, J.N.; project administration, D.S.L.; funding acquisition, J.N. All authors have read and agreed to the published version of the manuscript.

